# The Long-run Effects of Secondary School Track Assignment

**DOI:** 10.1101/599605

**Authors:** Lex Borghans, Ron Diris, Wendy Smits, Jannes de Vries

## Abstract

This study analyzes the long-run effects of secondary school track assignment for students at the achievement margin. Theoretically, track assignment maximizes individual outcomes when thresholds between tracks are set at the level of the indifferent student, and any other thresholds would imply that students at or around the margin are better off by switching tracks. We exploit non-linearities in the probability of track assignment across achievement to empirically identify the effect of track assignment on educational attainment and wages of students in the Netherlands, who can be assigned to four different tracks. We find that attending higher tracks leads to increases in years of schooling by around 1.5 years for students at the lowest and the highest choice margin, and wage gains of around 15% and 5%, respectively. Results at the margin of the two medium tracks show no effect on educational attainment, and decreases in wage of around 12% from attending the higher track. The negative returns for the medium margin and the relatively low returns for the higher margin (compared to the required educational investments) are partly mediated by motivation and study choice.

## 1 Introduction

The grouping of students according to educational achievement is common across educational systems worldwide. The motivation for such practices is that different students may require different environments and different levels of instruction to optimally develop their skills. Anglo-Saxon countries generally address this by selecting students into ability groups for different school subjects, while many Continental European countries sort secondary school students into different tracks that each have their own distinct curriculum. The use of tracking is continuously debated in both the public and the academic domain. Early empirical studies in economics have typically focused on the efficiency and equity implications of tracking systems, or on the effect of the exact age at which tracking takes place; see, e.g., [1, 2, 3, 4, 5]. Less attention has been paid to the allocation process that sorts students into different tracks. Tracking involves, either explicitly or implicitly, the use of ability thresholds in order to sort students. Students above a particular threshold are deemed fit to attend the higher track, while those below are projected to be better off in a lower track. A crucial question is whether the exact location of such a threshold in the ability distribution is optimal for students that fall to either side of that threshold.

The aim of this paper is to estimate the long-run effects of track assignment for students who are at the achievement margin. The empirical analysis of this study is focused on the Dutch educational system, where track assignment is partly determined by scores on an achievement test taken at the end of primary school (grade 6). This test contains certain threshold scores, which indicate the required ability level for a specific track. In reality, the adherence to these threshold scores by schools is lenient, but they still induce non-linearities in the probability of treatment across achievement that are not proportional to differences in the ability and potential of these students. This can be exploited to identify the effects of track assignment on future outcomes. We match survey data on Dutch secondary education students with administrative data on educational attainment and job market information in later life. Job market information is available up to an age of 42 years old. The Dutch educational system contains four different tracks in the period under analysis, which implies that there are three choice margins for which the effect of track assignment can be estimated. We find that attending the higher track increases educational attainment by around 1.5 years for the lowest and the highest choice margin. The subsequent labor market returns from attending the higher track are around 15% in the former and around 3-7% in the latter case. In contrast, we find that attending the higher track at the medium margin has no effect on years of schooling, and lowers wages by around 12% in the long run. These effects are partly mediated by study choice and motivation.

The choice of which track to attend typically involves schools, students and parents. Parental aspirations generally lean towards higher/more academic tracks that involve better peer quality and provide more direct paths towards high levels of post-secondary education [6]. Social status considerations and overconfidence could lead students to attend tracks that are too demanding and thereby hinder learning; see, e.g., [7] for evidence on education as a positional good and [8] for evidence on overconfidence in self-assessment. [9] find that a shift from parental to teacher influence for track assignment in Germany has led to a reduction in grade retention in secondary school, providing suggestive evidence that parents are indeed prone to push their children into too demanding tracks. Conformity to the educational level of the parents could make especially those with highly educated parents prone to attend too high tracks (and those with low educated parents to attend too low tracks). Schools face other considerations when setting thresholds for track attendance. Lower entrance requirements attract more students, while higher entrance requirements improve average peer quality within each track, and decrease the risk of costly grade retention. Hence, there are several reasons why the achievement threshold is not set at the indifferent student.

Research on tracking has traditionally focused on estimating effects for school achievement and educational attainment. To assess the effectiveness of tracking or track assignment, it is especially important to look at how tracking affects labor market outcomes, as that is ultimately what students are prepared for in these tracks. In fact, an increase in the number of completed years of schooling represents a cost from an economic perspective, and can only be beneficial when such an investment produces positive returns in meaningful later-life outcomes. While average wage returns to extra schooling are high, they are also strongly heterogeneous [10]. Increasing the size of the higher track can lead to increases in educational attainment simply because more students are eligible for higher levels of post-secondary education. Additionally, shifting from an academic to a vocational focus might lead to decreases in academic achievement, but could be to the benefit of other skills. Recent studies that estimate long-run effects underline this. [11] show that a tracking reform in Romania led to an increase in the number of students that completed an academic track, but not to increases in labor market outcomes. Similarly, [12] analyzes a policy change in Sweden that gave the vocational track a more academic curriculum and identifies an increase in educational attainment in secondary school, but no effect on earnings. Additionally, [13] show that the life-cycle dynamics of students following vocational tracks and students following academic tracks are different, underlining the importance of measuring wage effects at multiple ages across the life cycle.

These studies analyze policy changes that changed the content of tracks. In contrast, we analyze assignment of students to tracks for a given set and content of tracks. As such, this paper relates most closely to a recent study by [14], who estimate the long-run effect of track attendance by exploiting the fact that relatively younger students are less likely to attend higher tracks because of the month of birth effect. Our study estimates a similar effect, but at a different margin. [14] identify a local average treatment effect (LATE) for those that would attend a higher track if they would be born earlier in the year. We elicit a different local effect, namely for those who would attend a higher track if their achievement would have been marginally higher. This particular LATE reflects a local effect that is critical for decision-making with respect to tracking. It answers whether ability thresholds are indeed set at the indifferent student or whether students around the margin would be better off by switching to another track. Additionally, our study adds to the literature by estimating the effects of track attendance at multiple margins, and for multiple cohorts in time.

The organization of this paper is as follows. Section 2 specifies the theoretical framework of this study. Section 3 gives a brief overview of relevant characteristics of the Dutch educational system. Section 4 discusses the data. Section 4.2 presents the methodology, while main results are discussed in Section 5. Robustness analyses are presented in Section 6. Section 7 concludes.

## 2 Theory

Countries differ in how they assign students to tracks, but track assignment generally relies largely on measures of student achievement. This can rely on measures of general ability or on how students perform by type of subject (e.g. in quantitative subjects versus languages). Additionally, in countries in which tracks have strongly differentiated curricula, student preferences are a leading determinant. In our empirical setting, track assignment is based on a measure of overall ability, which is why the theoretical framework is constructed from this perspective as well.

We define an overall ability indicator *θ_i_*, on which track allocation decisions are based. Future outcomes (*Y_i_*) for students (e.g. future wages) depend on *θ_i_* and the track T the student attends. We specify a simple linear relation:

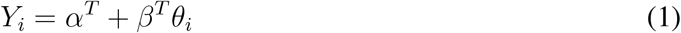

In the remainder of this section we assume, without loss of generality, that there are three tracks: low (L), medium (M) and high (H). Each track has its own curriculum (including not only the set of courses, but also level and pace of instruction). The level of these curricula are geared towards the average ability of students in that track. As such, lower tracks produce more favorable outcomes for low-ability students and higher tracks for high-ability students. In our linear framework, this implies that *α^L^*>*α^M^*>*α^H^* and *β^L^*<*β^M^*<*β^H^*. This situation is depicted on the left side of Fig 1. In the figure, it is assumed that allocation of students to tracks is efficient: no student can switch and make themselves better off. In other words, outcomes equal:

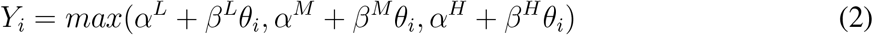

**Fig 1.**
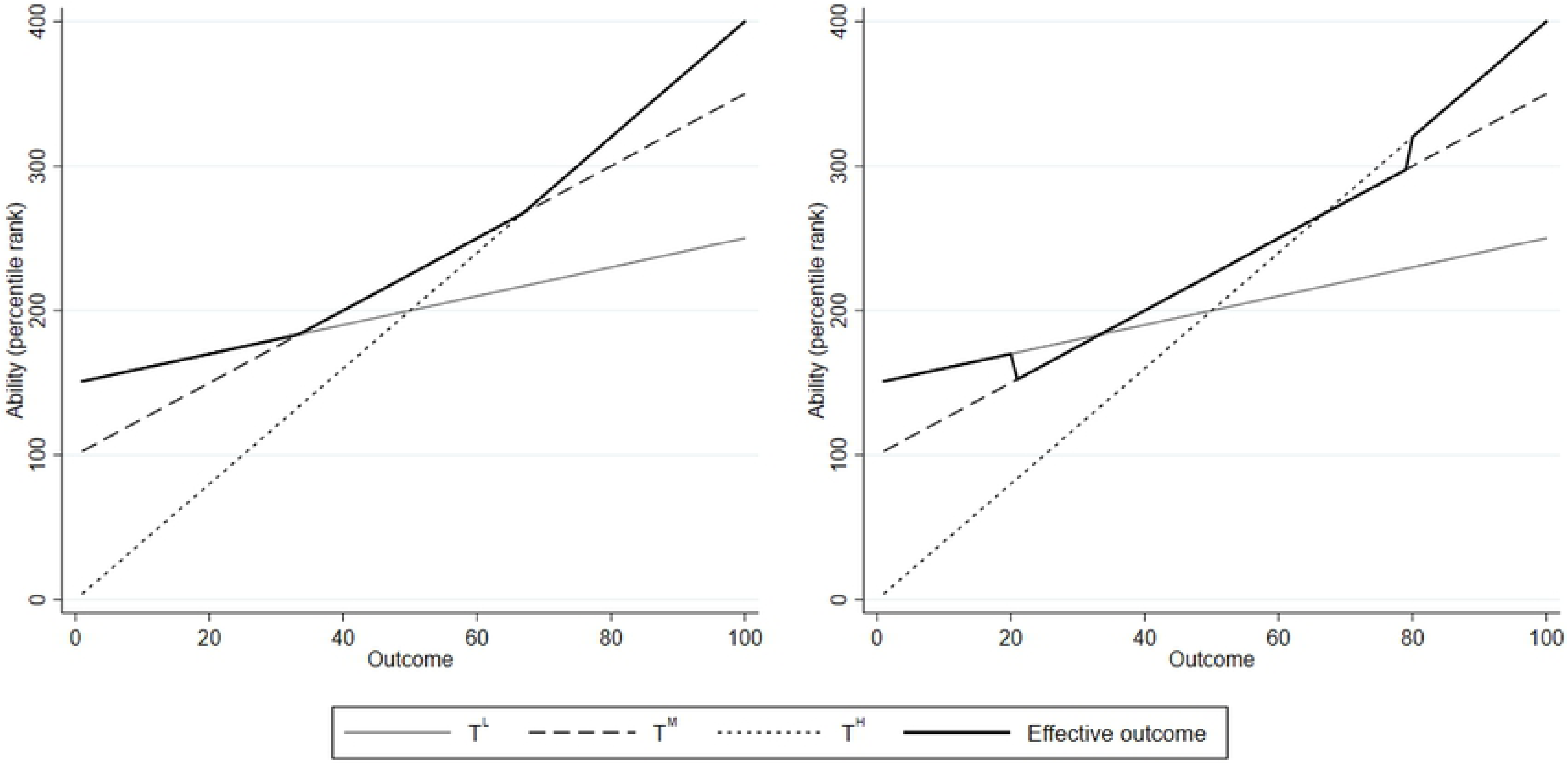
Allocation of students to tracks: theoretical framework. **Note:** The figure shows a theoretical depiction of the relation between outcome *Y* and attendance of low (L), medium (M) and high (H) tracks, across the ability distribution. On the left side, track assignment is efficient and the effective outcome line traces out the maximum outcomes. On the right side, the lower threshold is more strict and the higher threshold is more lenient, leading to discontinuities in outcomes.

In this scenario, the threshold ability levels lie before the first student in the distribution for which the higher track is the optimal choice and after the last student in the distribution for which the lower track is the optimal choice. When thresholds are located at different points, some students can be better off by switching. The effective outcome line will not follow the highest points and discontinuities in the outcome variable across the distribution will appear. The right side of Fig 1 describes the cases where the lower threshold is more lenient and the higher threshold is more strict. In the former case, the lowest-ranked students in the medium track would be better off in the low track and hence there is a downward jump between the highest low-track students and the lowest medium-track students. In the latter case, the highest ranked medium-track students would be better off in the higher track and there is an upward jump in the effective outcome line.

In reality, *θ_i_* is not directly observed and typically proxied with (noisy) measures of school achievement. This also applies to the empirical setting of this paper. Using a noisy ability signal for student sorting automatically implies that allocation is not fully efficient and some students would be better off in another track than where they were assigned to. Given that noisy ability signal, outcomes are still maximized when the threshold achievement level is set at the indifferent student (assuming noise is symmetrically distributed). In that case, *average* outcome lines would still be equal to the situation in Fig 1, and the average effective outcome line is smooth across the distribution. Put differently, efficiency gains can be made by using less noisy achievement measures or by putting thresholds at a more optimal position. Our focus in this paper is on the latter.

We do not consider peer effects explicitly in this model, which are incorporated within *α^T^* and *β^T^*. If one would assume a constant positive effect of better peers, *α^L^* would decrease (i.e. Track L is shifted downward) and *α^H^* would increase (i.e. Track H is shifted upward). Hence, peer effects reduce the part of the distribution for which the lower track is more optimal and increase the part of the distribution for which the higher track is more optimal. Explicitly modeling peer effects in the framework, however, provides no added value in the context of this study.

Additionally, we explicitly take an individual point of view. ‘Efficient’ assignment as in Fig 1 means that individuals cannot switch and be better off. The optimal allocation of students to tracks from a social welfare perspective represents a separate policy question. Changing effective thresholds also changes peer quality, pace of instruction etc. This would lead to changes in the parameters *α^T^* and *β^T^*. If one assumes that all students benefit from higher average peer ability then thresholds that maximize individual outcomes given the assignment of the rest of the distribution (as in Fig 1) is too lenient from a social welfare perspective, ceteris paribus (more so if we also consider general equilibrium effects). Stricter thresholds would be more optimal since they increase peer quality in tracks on both side of the threshold. Nonetheless, the framework as depicted is valuable as it identifies gains and losses for individuals who are at the choice margin for two specific tracks, which represents crucial information for students and their parents who face this choice. When we refer to ‘optimal’ track allocation in the remainder of this paper, this pertains to this individual perspective and not to a social optimum.

As argued in the introduction, the impact of track assignment can potentially differ between different outcome measures. In the context of the theoretical framework, this means that *α^T^* and *β^T^* are dependent on the defined outcome variable. For example, positive impacts on educational attainment could arise even in the absence of better student learning, because higher tracks make students eligible for higher levels of post-secondary education. Although literature clearly shows that the average return to an extra year of schooling is positive and substantial, this return can be different for students who are induced to prolong their educational career because of more lenient requirements. This underlines the value of measuring also the labor market effects of such treatments.

## 3 Dutch educational system

The Dutch educational system is characterized by relatively early tracking and a high number of tracks. Fig 2 provides a schematic overview from primary to tertiary education. Primary education lasts six years (preceded by two years of kindergarten). The focus of this study is on tracking in secondary education, and on how this influences post-secondary trajectories.

**Fig 2.**
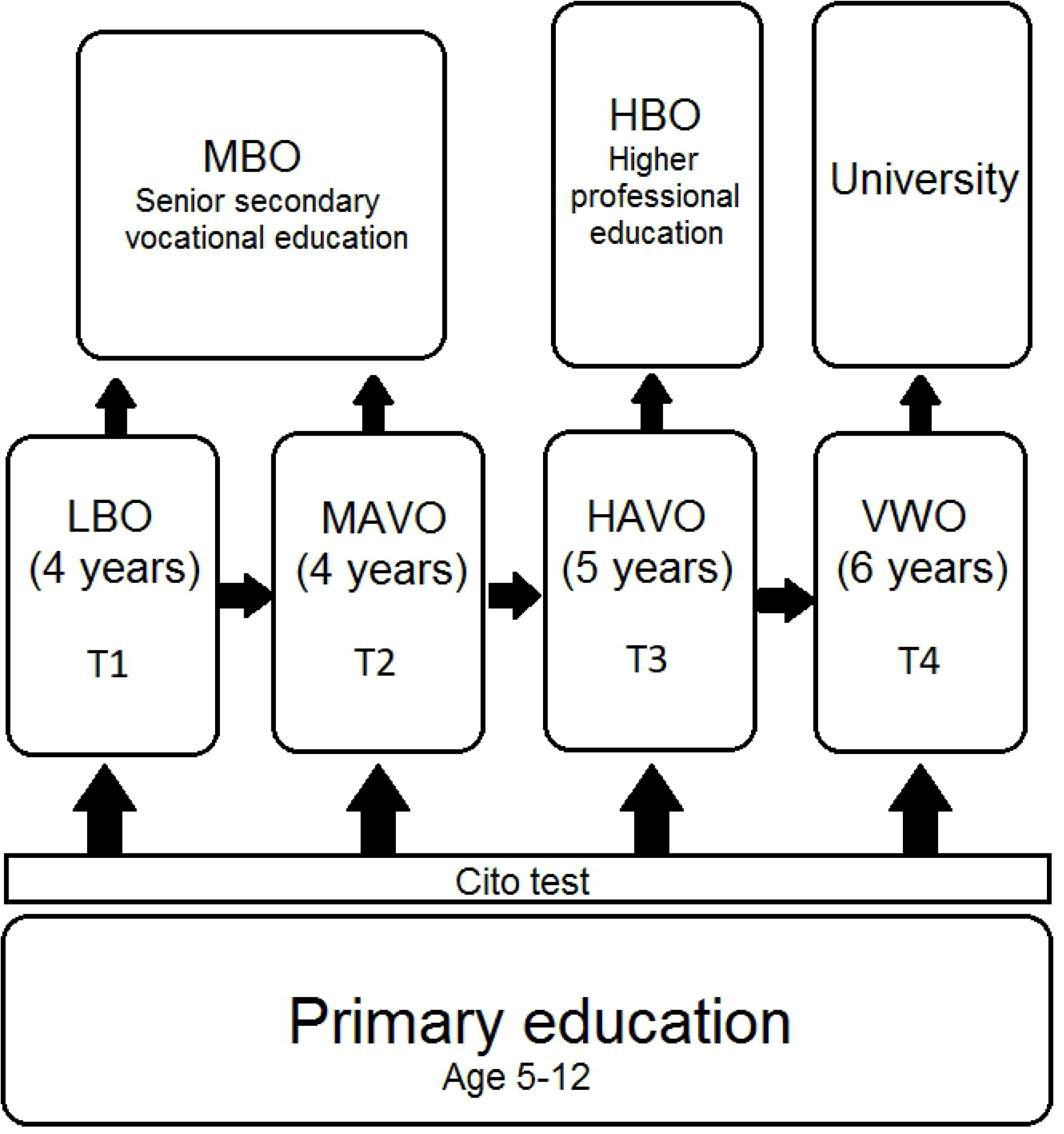
Dutch educational system. Source: Center of International Education Benchmarking.

### 3.1 Secondary education

After finishing primary school, students in the Netherlands are selected into four main tracks: *lbo* (vocational), *mavo* (lower general), *havo* (higher general) and *vwo* (pre-university). Students can be relegated to lower tracks after track assignment, in case of low achievement, but moving to higher tracks is generally only possible when the current track has been completed. The *lbo* and *mavo* tracks have been merged in 1999 into the joint *vmbo* track. Within *vmbo*, there still exists a practical and a theoretical subtrack. For the vast majority of students in our sample, the system with four tracks applies. For the remainder of this paper, we refer to the available tracks as T1 for *lbo*, T2 for *mavo*, T3 for *havo* and T4 for *vwo*.

A leading determinant of track assignment is the 6th grade exit test that primary schools are obliged to administer. While several alternative tests exist, a large majority of 85% of Dutch primary schools administer the standardized ‘Cito test’ [15]. The obtained Cito score is connected to a ‘teacher recommendation’ for any of the four tracks, which is given by the 6th grade teacher and sent to the prospective secondary school. Recommendations can be mixed, when students’ achievement level is around the margin of the required level for a certain track. Students and their parents can decide to deviate from the track recommendation, if secondary schools allow for this. The latter are obliged by law to consider at least one of the two sorting mechanisms (test score and/or teacher recommendation) when admitting or sorting students. However, secondary schools are free to decide their exact assignment rules, and to deviate from the score thresholds that the manufacturers of the Cito test report.

There is an additional opportunity for students to be selected into a track that does not correspond to their Cito score, because final selection can be postponed until the second or third year of secondary education. This occurs through the existence of temporary comprehensive grades (so-called ‘bridge-grades’), where students of two or more tracks are still kept together. This is most common for Track 3 and Track 4 students, while students in the lower two tracks are generally selected early. In our sample, roughly 90% of T3 and T4 students is in a comprehensive grade for at least one year, and 35% for 2 years. About 75% of students in Tracks 1 and 2 is in a specific track in the first year of secondary education already. When track assignment is postponed to later years, it is also based on student achievement, most commonly through school-specific requirements with respect to grade point averages.

Hence, while achievement is the key driver of track sorting in the Netherlands, these institutional features imply that assignment is far from deterministic and that there is considerable leeway in getting into tracks that do not match up with ‘eligibility’ status of students if we would follow only the achievement test. We elaborate on the patterns of student sorting by achievement and their implications for the empirical approach in Section 4.2.

### 3.2 Post-secondary education

The Dutch educational system has three levels of post-secondary education. The lowest level is *mbo*, which has a vocational orientation. Higher education can be divided into two categories. *Hbo* provides higher professional education (also known as vocational university), while *wo* consists of university education. Students with a high school diploma from the T3 track or higher can enter *hbo*, while *wo* is only available for students who complete T4. Completing the highest level of *mbo* makes one eligible for entering *hbo* as well, while completing the first year of *hbo* gives direct access to university education. Hence, it is still possible for students from lower tracks to complete higher education, although the route is less direct and involves additional time. Nonetheless, around 15% of students from the T2 track in our data sample still complete higher education. For the most recent cohorts, this even equals 22% (see Appendix Figure A1).

## 4 Materials and Methods

### 4.1 Data

For the empirical analysis, we link several data sources. Secondary school data are collected from the Secondary Education Cohort Studies. These are large representative longitudinal surveys of Dutch pupils whose educational career was followed from the first year of secondary education (around age 12) until they leave full-time education. They are carried out by Statistics Netherlands and the Groningen Institute for Educational Research; see, e.g., [16, 17]. These data include measures of student achievement, student background, and the attended level of education for each year. We use cohort studies starting in 1977, 1983, 1989, 1993 and 1999.

All students that participated in the Secondary Education Cohort Studies are matched to administrative information from the System of Social Statistical Datasets (SSB) on educational attainment, wages and labor market status. The main outcome measure for educational attainment is years of schooling, which is coded towards the highest obtained degree, and is available until the year 2008. The labor market data contain the monthly earnings that are reported to the tax authorities, in every year from 2001 to 2007. These earnings are registered for the month of September in that particular year. The earnings for this month are typically seen as most representative, as they are generally not affected by end-of-year bonuses or vacation pay.

For the empirical analysis, we recode this variable. As we are mainly interested in wage effects and not in labour supply effects, we correct for the full-time equivalent of the job (number of hours worked per week divided by the number of hours for a full-time worker with this particular job). Additionally, we exclude (corrected) monthly wages below 1300 euro. This approximates the minimum wage in the Netherlands in this period, and values below this threshold suggest that the individual was not employed for the full month. Finally, we topcode at 20,000 euro per month to mitigate the influence of high-earning outliers. Our main outcome variable averages this recoded indicator across the seven years for which data are available. We refer to this indicator as the ‘mean wage’, given the correction for hours worked. Section 5 discusses the difference between the main estimates and those based on raw earnings.

The matched cohorts comprise around 25,000 observation for the 1977 cohort and around 15,000 observations for later cohorts. Labor market information is available from those who just entered the labor market until those who are 42 years old (the 1977 sample in 2007). The 1999 dataset includes individuals who are generally too young to have entered the labor market in any of the years for which we have data, hence long-run outcomes are not estimated for this cohort. Labor market effects for the 1993 cohort are not estimated for methodological reasons (see Section 4.2). Around 10% of students in the 1993 cohort is still in education by 2008. Robustness checks show that the results are not sensitive to this data limitation: we obtain virtually identical results when we assume that all these students finish the study they are currently attending. The share of students still in education is negligible for all other cohorts.

The 1989, 1993 and 1999 cohorts also contain data on 9th grade achievement. The test scores in the 1989 and 1993 cohorts suffer from a high number of missing observations. We find that the probability of taking the test is positively related to previous achievement of the student (i.e. the better students in class are more likely to take the test). This can severely confound the estimates of our analysis, since the empirical approach effectively compares students at the margin. For these two cohorts, we use two imputation approaches that assess sensitivity to the missing value issue. For the 1999 cohort, the number of missing observations is negligible. Grade 9 test scores are available for language, mathematics and general problem-solving.

Unfortunately, the specific results of the Cito test that largely determines track assignment is only available for the 1999 cohort. For the other cohorts, a very similar test is available which is taken at the start of secondary education. This test is based on the same pool of questions and therefore serves as a proxy for the actual Cito test. This test is labeled as the *Entrance Test*, since it is taken just after entering secondary education. Entrance Tests are exactly identical for the 1989, 1993 and 1999 cohorts and contain 20 questions each in math, language and information processing. The test for the 1983 cohort contains the same division, but with different questions, while the 1977 Entrance Test contains 45 questions in language and 25 questions in math. Additionally, students from the 1983 cohort already take the test at the end of grade 6.

Summary statistics are provided in Table 1. The share of students attending T1 slightly increases over the cohorts, mainly at the expense of T2. As in other developed countries, we observse a steady increase in obtained years of schooling over time. Parental education levels are increasing as well across cohorts, reflecting that this upward trend was already present in earlier cohorts. There is some variation in the mean levels of other background variables, but no clear trend nor marked differences in the background of students across these cohorts.

**Table 1:** Summary statistics

|  | 1977 |  | 1983 |  | 1989 |  | 1993 |  |
| --- | --- | --- | --- | --- | --- | --- | --- | --- |
|  | Mean | Std. dev. | Mean | Std. dev. | Mean | Std. dev. | Mean | Std. dev. |
| Track 1 | 0.256 | 0.436 | 0.323 | 0.468 | 0.333 | 0.471 | 0.359 | 0.480 |
| Track 2 | 0.422 | 0.494 | 0.367 | 0.482 | 0.381 | 0.486 | 0.323 | 0.468 |
| Track 3 | 0.140 | 0.347 | 0.142 | 0.349 | 0.135 | 0.341 | 0.152 | 0.359 |
| Track 4 | 0.182 | 0.386 | 0.168 | 0.374 | 0.151 | 0.358 | 0.165 | 0.371 |
| Entrance Test | 42.17 | 12.11 | 33.32 | 10.17 | 34.23 | 11.33 | 35.20 | 11.42 |
| Years of schooling | 12.27 | 3.29 | 12.64 | 3.54 | 13.05 | 3.83 | 12.99 | 3.84 |
| Wage 2007 | 3765.85 | 2163.48 | 3452.53 | 1709.74 | 2878.54 | 1117.40 | 2460.09 | 874.17 |
| Wage 2006 | 3669.68 | 2109.74 | 3300.06 | 1628.58 | 2705.51 | 1066.36 | 2271.88 | 834.21 |
| Wage 2005 | 3290.86 | 1817.53 | 2905.21 | 1264.19 | 2428.57 | 836.90 | 2045.64 | 752.97 |
| Wage 2004 | 3186.34 | 1680.56 | 2793.78 | 1151.76 | 2322.09 | 753.22 | 1917.74 | 688.77 |
| Wage 2003 | 3103.36 | 1576.27 | 2696.45 | 1076.33 | 2213.90 | 704.18 | 1763.19 | 652.08 |
| Wage 2002 | 2977.85 | 1488.49 | 2568.05 | 1026.50 | 2047.57 | 660.85 | 1544.12 | 628.42 |
| Wage 2001 | 2832.72 | 1412.19 | 2418.05 | 927.00 | 1872.69 | 647.18 | 1312.55 | 604.68 |
| Female | 0.503 | 0.500 | 0.504 | 0.500 | 0.481 | 0.500 | 0.485 | 0.500 |
| Big 4 cities | 0.091 | 0.288 | 0.055 | 0.228 | 0.045 | 0.207 | 0.060 | 0.238 |
| Non-Dutch | 0.079 | 0.265 | 0.086 | 0.275 | 0.109 | 0.311 | 0.090 | 0.286 |
| High parental educ. | 0.164 | 0.370 | 0.147 | 0.354 | 0.195 | 0.396 | 0.220 | 0.414 |
| Low parental educ. | 0.211 | 0.408 | 0.329 | 0.470 | 0.153 | 0.360 | 0.086 | 0.281 |
| Low social class | 0.120 | 0.325 | 0.111 | 0.314 | 0.087 | 0.282 | 0.085 | 0.279 |
| High social class | 0.115 | 0.318 | 0.107 | 0.310 | 0.145 | 0.352 | 0.136 | 0.343 |

**Notes:** The table shows means and standard deviations of the main variables, separately for each cohort. Wages are in euro's, per month, pre-tax, corrected for full-time equivalent of the job and topcoded at 20,000. The variables Track 1 to Track 4 report the fraction of students that are assigned to that track. The Entrance Test includes 70 questions in 1977 and 60 questions in all other cohorts. Years of schooling is coded to the highest degree obtained. 'Big 4 cities' indicates whether the child lives in one of the four big cities in the Netherlands.

### 4.2 Methodology

#### 4.2.1 Non-linearity design

The aim of this study is to estimate the effect of track assignment on educational attainment and wages. As specified in the theoretical framework, one can find the effect of track assignment by identifying discontinuities in outcome variables around the achievement margin between two specific tracks. As such, the setup would be well-suited for a regression discontinuity design (RDD), which exploits discontinuities in treatment at a specific threshold value of a forcing variable. This approach is, however, empirically not feasible here. For one, we only observe the true forcing variable in one of the cohorts (1999). More importantly, the variability in the thresholds that schools adhere to and the use of postponed tracking lead to a very high fuzziness of track allocation around the threshold. Figs 3-5 show the relation between treatment probability and the Entrance test, for all three margins (Fig 4 excludes T4 students, as we need the relation between achievement and track assignment to be monotonic; as shown later, including only the lower three tracks is the preferred specification in the empirical analysis). The figures consistently show that a true discontinuity in treatment is absent. Moreover, Appendix Figure A2 shows that in the cohort where we do observe the official Cito score, the pattern of treatment probability across score is similar to that for the Entrance Test. Hence, the lack of a strong discontinuity in treatment at or around the achievement threshold is not driven by the use of the proxy test, but by the specific dynamics of student sorting in the Dutch system.

**Fig 3.**
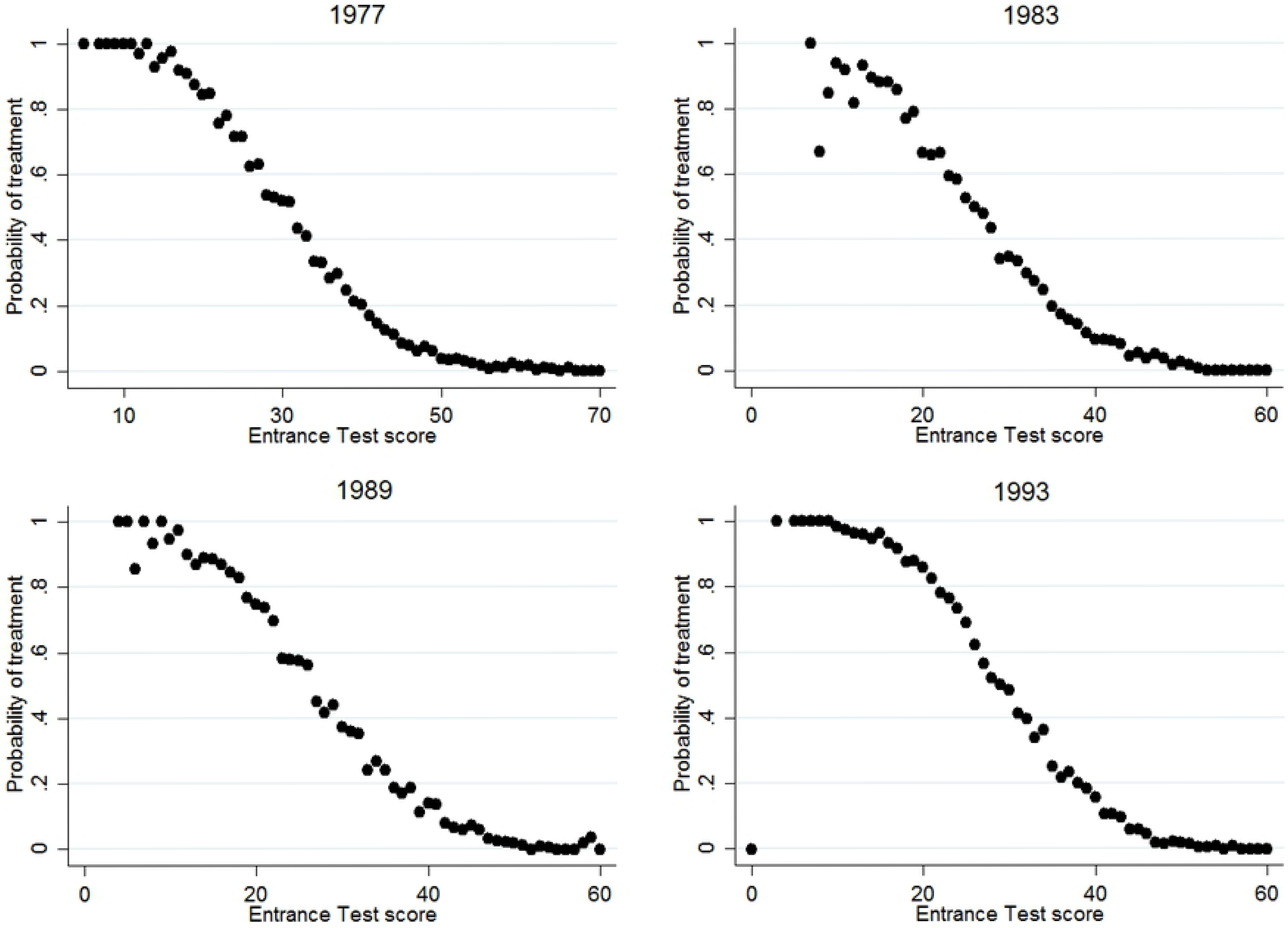
Share of students assigned to T1, across test scores and cohorts. **Note:** The figure shows the share of students that are assigned to the T1 track, for every score on the Entrance test, separately for all cohorts.

**Fig 4.**
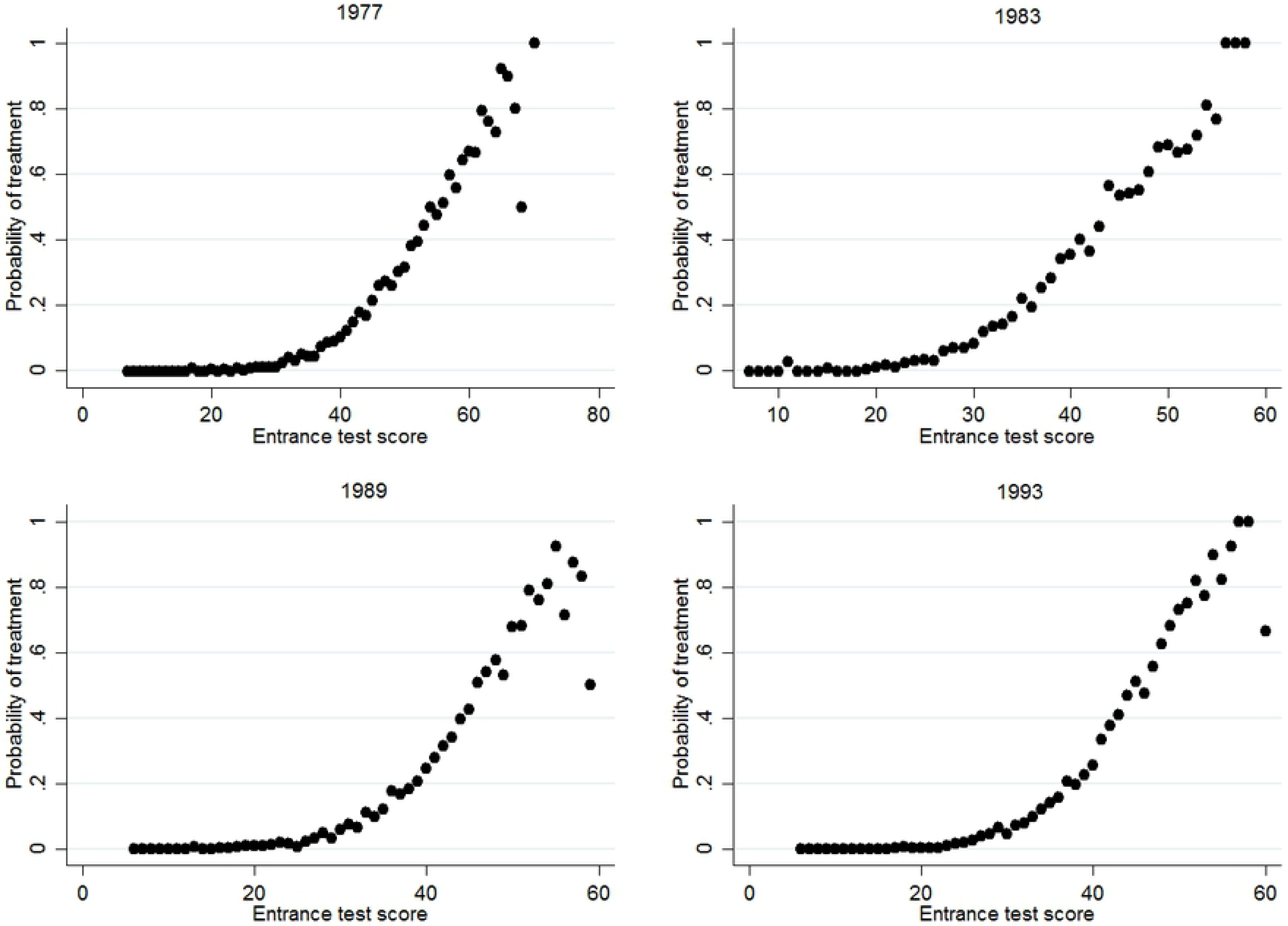
Share of students assigned to T3, across test scores and cohorts. **Note:** The figure shows the share of students that are assigned to the T3 track, for every score on the Entrance test, separately for all cohorts. Students that attend T4 are excluded in the calculation.

**Fig 5.**
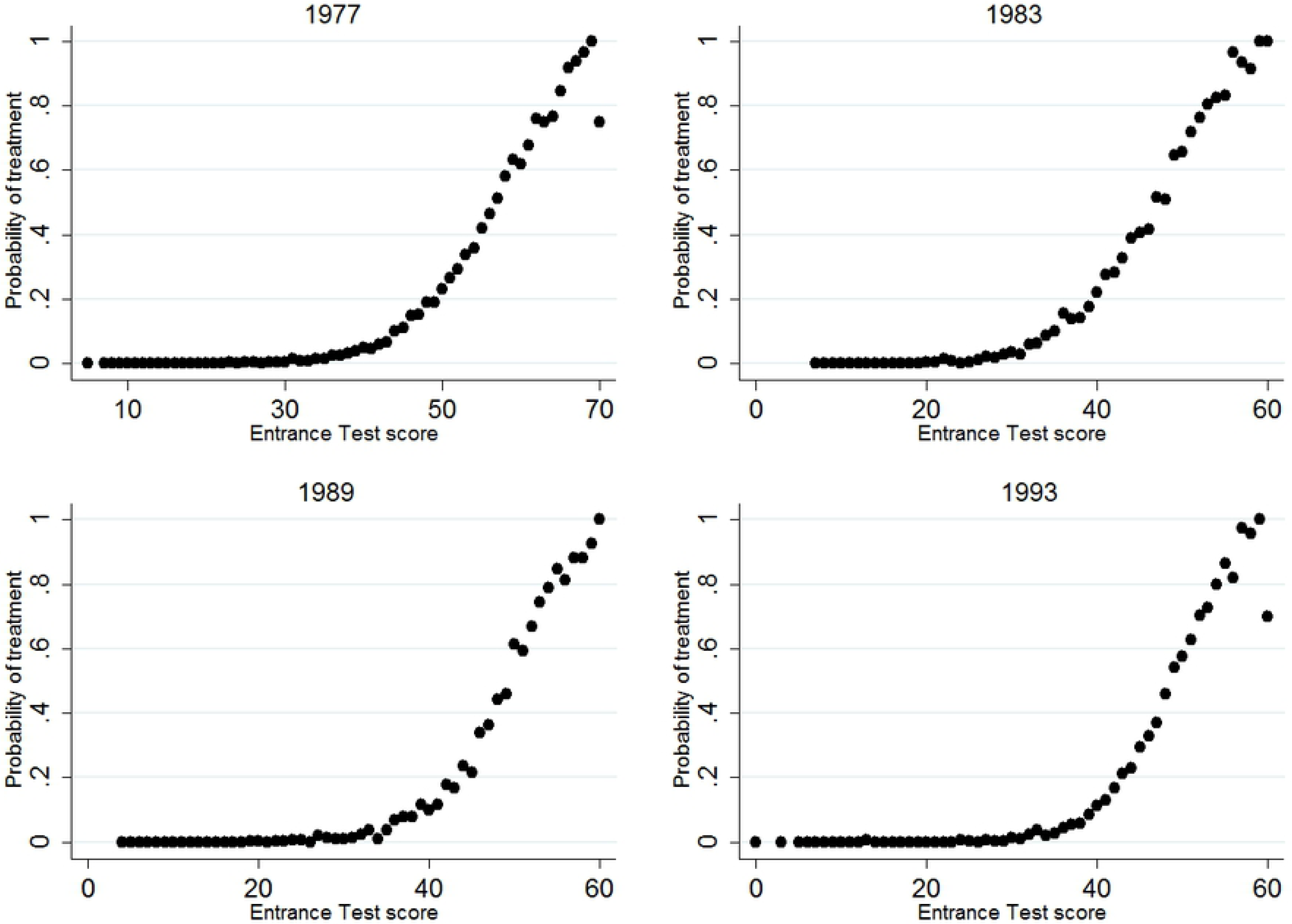
Share of students assigned to T4, across test scores and cohorts. **Note:** The figure shows the share of students that are assigned to the T4 track, for every score on the Entrance test, separately for all cohorts.

Although Figs 3-5 fail to show a strong discontinuity in treatment probability, they do inhibit a non-linear pattern that can potentially be exploited. Because track assignment is based on achievement, the probability of being assigned to a certain track is highly responsive to changes in achievement around a certain (implicit) threshold score. The treatment probability increases strongly around this score, while it remains flat in segments before and after. In other words, the increase in the probability of treatment is disproportionally low for increases in score that are far from the threshold and disproportionally high for increases in score around the threshold. This ‘disproportionality’ can be exploited to estimate the effect of track assignment, through a two-stage model in which the fraction of students that is treated for a given test score acts as an instrument for the treatment indicator. Such a conditional mean approach does not rely on any defined threshold score, as it incorporates all changes in probability within the defined bandwidth.

A first condition for applying this approach is that the relationship between the outcome variable and the forcing variable has a different functional form (net of treatment) than the relationship between the treatment variable and the forcing variable. Fig 6 displays the relation between educational attainment or wages and the Entrance Test score, for the sample as a whole. Appendix figure A3 shows the same relation split across tracks. These figures show a linear pattern for educational attainment in all cohorts and for wages in the three oldest cohorts. The pattern for the 1993 cohort is strongly non-linear as many high achieving students have not entered the labor market yet, leading to two opposing forces in the relation between achievement and wages. Consequently, the optimal bandwidth is too narrow for a feasible first stage and wage effects are therefore not estimated for the 1993 cohort.

**Fig 6.**
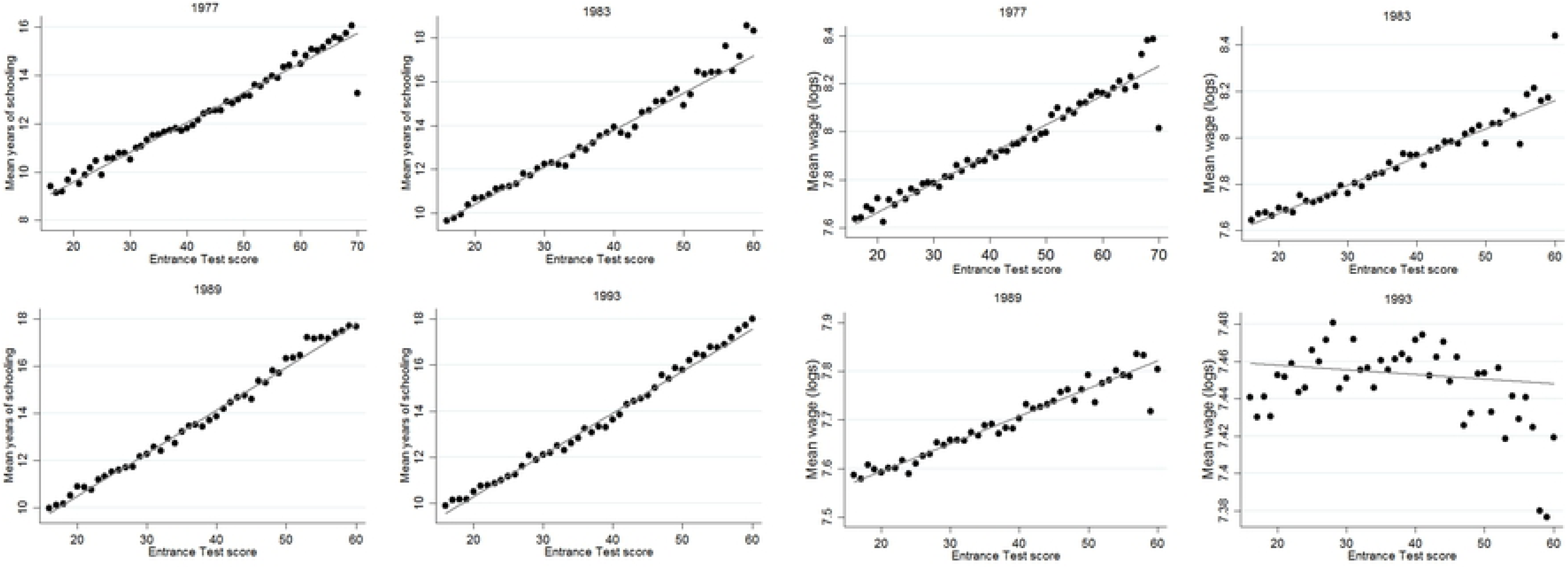
Mean years of schooling and wages across test scores and cohorts. **Note:** The figure shows the mean years of schooling (left half) and mean log wages (right half) for every score on the Entrance Test, separately for all cohorts.

A second condition is that there can be no similar non-linearities in other determinants of *Y_i_*, as these will be attributed to the treatment effect. Fig 7 shows the relation between our main outcome variables and an imputed vector of observable characteristics. This imputed vector combines information on gender, month of birth, whether the student lives in one of the four big cities in the Netherlands, ethnicity, and dummy variables for parental social status (six categories, based on occupation) and parental education (three categories). It is constructed by regressing the outcome (years of schooling or wages) on this set of controls and then fitting the predictive values. Hence, control variables that relate more strongly to the outcome receive a higher weight in the vector. The figures show that the relation between the imputed control vector and the main outcomes is highly linear the vector is constructed with respect to years of schooling, in all cohorts. With respect to wages, the patterns are also highly linear for the 1977 and 1989 cohorts. For the 1983 cohort, there is some degree of non-linearity at the very high end of the distribution, but the number of observations for these high scores is low. We assess sensitivity towards the specified bandwidth in Section 6, which will provide an indication to what extent this can bias results. The pattern for the 1983 cohort is also more erratic, as the test taken in that year is somewhat less predictive of future outcomes.

**Fig 7.**
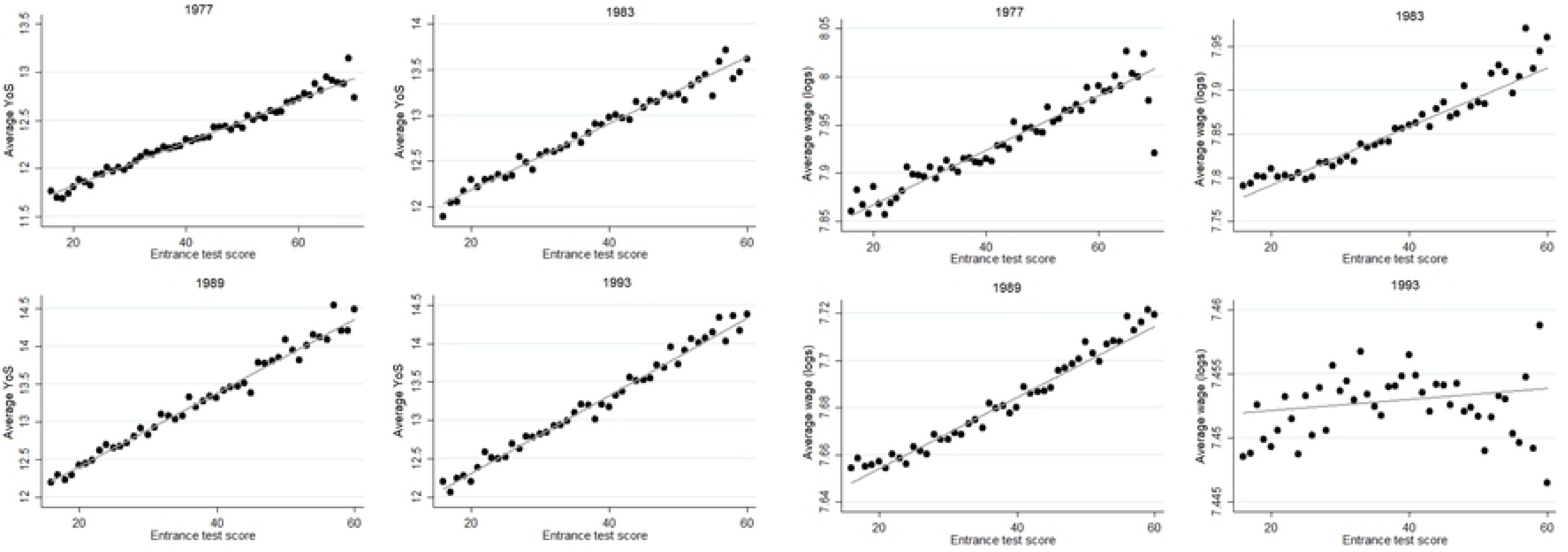
Control vector by Entrance Test score. **Note:** The figure shows the average values of a control vector for every score on the Entrance Test, separately for all cohorts. The control vector is constructed by regressing either years of schooling (left half) or log wage (right half) on a set of controls and then fitting the predictive values. Controls are: gender, month of birth, urbanization, ethnicity, dummies for parental social status (six categories) and dummies for parental education (three categories).

Appendix figures A4 to A7 show the relation to the Entrance Test for all controls separately. For the purpose of this exercise, parental education and parental occupation have been expressed as continuous variables rather than categorical dummies as in the estimation model (occupational categories are ranked by average wage). These separate relations also show highly linear patterns. Modest exceptions occur for gender in 1983 and urbanization in 1993. Note that the latter has comparatively modest explanatory power towards outcomes. The former may explain the non-linearity at the higher end for the 1983 cohort in Fig 7. We assess sensitivity to the inclusion of control variables in Section 5, which indicate to what extent these particular cases lead to sensitivity in the results. The strongly linear patterns between *S_i_* and the main outcome variables and between key observable characteristics and the main outcome variables in the large majority of cases provide evidence in favor of the validity of the assumption that the non-linearity in the relation with *S_i_* is exclusive to the probability of track assignment. Given a wide enough bandwidth, we can exploit this non-linearity to estimate the effects of track attendance.

The Two Stage Least Squares (2SLS) model becomes:

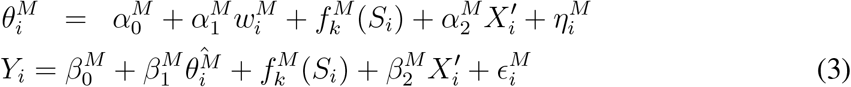

where *w_i_* = *E*[*θ_i_|S_i_*] (the conditional mean), which serves as the instrument for track attendance. The outcome *Y_i_* represents measures of either educational attainment or wages. For each outcome, three different treatment effects 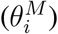 are estimated for each of the three margins *M*: T2 vs. T1; T3 vs. T2 and T4 vs. T3. The function *f_k_*(*S_i_*) represents the control function for the forcing variable, i.e. the Entrance Test. We employ a linear control function in all of the main analyses. We further include a vector of observable background characteristics (labeled *X*′). This is the same set of controls as used in Fig 7. Note that Model 3 is feasible, as long as 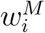 is a strong enough predictor of 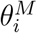, conditional on *S_i_* and 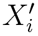, which would ensure a sufficiently strong first stage. For this to be the case, 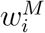 needs to have a different functional form in relation to 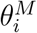 than the other variables in the model (within the specified bandwidth).

The approach we use bears similarities to that of [18], although in a different setting. They show that non-linearities in hedonic markets can be exploited in an IV approach through the calculation of conditional mean functions. The approach assumes that all unobserved determinants of the outcome are linearly related to *S_i_* across the bandwidth, and therefore captured by the control function. In other words, the model assumes that *∊* is mean independent of *S_i_* (*E*(*∊*|*S_i_* = 0)). This assumption is stronger than for an RD design, which requires that other (unobserved) determinants of the outcome are smoothly continuous at the threshold. On the other hand, our model relies on weaker assumptions than an OLS model, which requires that such determinants are completely unrelated to the treatment.

As mentioned before, part of the low adherence to threshold achievement levels occurs because different schools may employ different thresholds or different levels of strictness in adhering to thresholds. This may result in students that are just below the margin for one school to actively ‘shop’ for schools until they find a school that allows entry into the higher track. We emphasize that this behavior is not a threat to our identification, as long as our assumption of a linear relation between the outcome and unobservable determinants holds.

#### 4.2.2 Bandwidth selection

As our objective is to estimate track assignment effects at the achievement margin, we mainly want to select observations close to the (implicit) achievement margin. As with the RDD, we therefore need to establish bandwidths for our and assess whether the first stage of the model is still valid for the resulting subsample. We follow the typical approach used for RDD models by executing a cross-validation (CV) procedure to identify the bandwidth that minimizes the mean squared error of the control function [19]. The CV procedure weighs the additional power of including more observations against the loss in precision from moving further away from the threshold. As there is no explicit threshold score, we set the implicit threshold at the score where the treatment probability surpasses the 50% mark. We then step-wise extend the bandwidth from this point on each side of the threshold, and assess which particular score range minimizes the mean squared error.

We execute the cross-validation procedure for different sample compositions with respect to the number of included tracks and elicit the optimal one. While it is sufficient to include only those two tracks at the threshold, inclusion of additional tracks provides additional power and precision, which can be especially valuable when dealing with tracks for which test scores are concentrated within a small range. Although including additional tracks can lead to a bias if the relationship between *S_i_* and *Y_i_* strongly differs across tracks, the specifications that include additional tracks should be rejected by the CV-procedure in such a case in favor of specifications that only include the two tracks at the margin. In the sensitivity analysis, we address how sensitive the results are to the number of tracks that are included in the estimation.

The optimal specifications and bandwidths are separately estimated for each cohort and for each outcome variable, and evidently also for each treatment effect. Results suggest using the full set of tracks for estimation of the effect of T4 vs. T3 (with the exception of educational attainment in 1993, where the recommended specification includes only the top two tracks), and to include the lowest three tracks both for estimation of T2 vs. T1 and estimation of T3. vs. T2. The optimal bandwidths for all treatment effects and cohorts can be observed in Table 3. Appendix Table A1 shows the first stage results, applying each of these bandwidths. All estimates of first stage power are strongly above conventional thresholds (with the exception of the estimation with respect to wages in the 1993 cohort, as argued before).

The first stage is sufficiently strong in these cases because these bandwidths are rather wide, thereby ensuring that the relation between *S_i_* and *θ_i_* is still non-linear within the estimation window. These bandwidths are nonetheless optimal according to the CV exercise, which confirms that the (net) relation between *S_i_* and the outcomes is strongly linear, and that the linear control function still provides a very good fit also when moving further away from the implicit achievement threshold.

One may still question whether this conditional mean approach estimates treatment effects at the margin, especially when also tracks that are not part of the choice margin are included in the estimation. The IV model estimates a LATE for students for which incremental changes in measured achievement induce changes in track attendance. Hence, segments of the distribution where the treatment probability remains flat in Figs 3-5 have no weight in the estimation of treatment effects. At the same time, the model puts especially strong weight on students that are at the (implicit) achievement threshold, where the slope in Figs 3-5 is steepest. Still, there might be heterogeneity in treatment effects within the increasing parts of the figures, and the estimates can as such be partly driven by the potentially differential treatment effects of those who are relatively further from the margin. We assess this concern by conducting a heterogeneity analysis (Section 5.4) and by inducing variation in sample composition (Section 6.3). Moreover, Section 6.3 will assess whether results still hold for more narrow bandwidths (with the caveat that very narrow bandwidths are not feasible in the model because the relation between *S_i_* and *θ_i_* would become linear).

## 5 Results

We now present the estimated effects of track assignment at all three margins. The coefficients always represent the effect of attending the higher of the two tracks at the margin. Our main outcome variables are wages and educational attainment. We also add estimation of track assignment for school achievement in grade 9 (Section 5.3).

Table 2 reports results of each treatment on years of schooling and wages for a simple OLS model that regresses the outcome variable on track assignment and the score on the Entrance Test. These OLS estimates are presented for comparative purposes. A direct comparison with the IV estimates is not straightforward as IV elicits a LATE while OLS elicits an average treatment effect (ATE). OLS estimates are still valuable as they likely provide an upper bound of the ATE of attending the higher track at the margin, as one would expect that students in the higher track are better endowed in unobserved characteristics that are positively related to educational attainment and wages (also conditional on *S_i_* and *X^’^*). Additionally, a comparison of OLS results with and without controls for observable characteristics can provide an indication of how sensitive OLS models are to selection bias.

**Table 2:** OLS estimates of the long-run effect of track assignment

|  | T2 vs. T1 |  | T3 vs. T2 |  | T4 vs. T3 |  |
| --- | --- | --- | --- | --- | --- | --- |
|  | YoS | Wage | YoS | Wage | YoS | Wage |
| Panel A: No Controls |  |  |  |  |  |  |
| 1977 cohort | 1.22***<br>(0.087) | 0.082***<br>(0.012) | 0.841***<br>(0.071) | 0.116***<br>(0.0094) | 1.61***<br>(0.080) | 0.188***<br>(0.0092) |
| 1983 cohort | 1.66***<br>(0.071) | 0.067***<br>(0.0075) | 1.18***<br>(0.103) | 0.101***<br>(0.010) | 2.27***<br>(0.099) | 0.162***<br>(0.0094) |
| 1989 cohort | 1.77***<br>(0.108) | 0.0082<br>(0.0081) | 1.80***<br>(0.115) | 0.060***<br>(0.0069) | 2.46***<br>(0.102) | 0.080***<br>(0.0072) |
| 1993 cohort | 1.81***<br>(0.099) | - | 1.85***<br>(0.102) | - | 1.61***<br>(0.107) | - |
| Panel B: With controls |  |  |  |  |  |  |
| 1977 cohort | 1.13***<br>(0.080) | 0.093***<br>(0.0084) | 0.776***<br>(0.071) | 0.109***<br>(0.0083) | 1.51***<br>(0.080) | 0.170***<br>(0.0083) |
| 1983 cohort | 1.55***<br>(0.072) | 0.096***<br>(0.0066) | 1.05***<br>(0.101) | 0.109***<br>(0.0093) | 2.11***<br>(0.100) | 0.153***<br>(0.0090) |
| 1989 cohort | 1.43***<br>(0.097) | 0.022***<br>(0.0066) | 1.53***<br>(0.106) | 0.063***<br>(0.0068) | 2.12***<br>(0.098) | 0.079***<br>(0.0074) |
| 1993 cohort | 1.56***<br>(0.094) | - | 1.60***<br>(0.100) | - | 1.48***<br>(0.100) | - |

**Notes:** \*Significant at 10% level \*\*Significant at 5% level \*\*\*Significant at 1% level

Table 2 shows that OLS estimates are consistently in favor of attending the higher track. Estimates are relatively highest for T4 vs. T3 treatment, for both outcomes. Coefficients indeed reduce when adding controls when using years of schooling as outcome, but this is less clear for the wage regressions. Wage estimates become even larger at the lowest margin when controls are added. This is, however, solely due to the control for gender (i.e. women are more likely to attend T2 vs. T1 for a given score, while they earn lower wages). As expected, those attending the higher track at the margin have higher educated parents from higher social classes, and controlling for these variables lowers treatment effects in the OLS model in all cases.

### 5.1 Educational attainment

The estimates of the long-run effects of tracking for the main IV model are presented in Table 3. Columns 1, 3 and 5 show how track assignment affects educational attainment, for all three margins. Higher tracks consistently lead to more years of education at both the lowest and the highest margin. The point estimates center around 1.5 extra years of education in each case. For T2. vs T1 treatment, the imprecision of the estimates is relatively high, and they are just shy of statistical significance in 1983 and 1989, but effect sizes are rather consistent. For T4 vs. T3 treatment, effects gradually increase by cohort, which could be related to the fact that higher education attendance has increased markedly over the same period. All estimates for the effect of T3 vs. T2 are statistically insignificant. Standard errors are large, especially for the 1983 and 1989 cohorts, but the point estimates are consistently low. Hence, it appears that attending a higher track for students at this specific margin does not translate into higher educational attainment. Additional analysis (not shown) indicates that students that attend T3 over T2 at the margin still predominantly complete Track 3 and also obtain slightly more *hbo* diplomas, but are simultaneously less likely to complete any post-secondary education.

**Table 3:** IV estimates of the long-run effect of track assignment

|  | T2 vs. T1 |  | T3 vs. T2 |  | T4 vs. T3 |  |
| --- | --- | --- | --- | --- | --- | --- |
|  | YoS | Wage | YoS | Wage | YoS | Wage |
| 1977 cohort | 1.76**<br>(0.706)<br>[10-47] | 0.147***<br>(0.039)<br>[12-59] | 0.204<br>(0.605)<br>[33-70] | -0.122***<br>(0.040)<br>[20-70] | 1.00***<br>(0.199)<br>[15-64] | 0.071***<br>(0.027)<br>[15-64] |
| 1983 cohort | 1.17<br>(0.753)<br>[22-48] | 0.152***<br>(0.051)<br>[15-55] | 0.010<br>(0.929)<br>[26-60] | -0.122<br>(0.086)<br>[25-60] | 1.25***<br>(0.339)<br>[21-60] | 0.027<br>(0.034)<br>[21-60] |
| 1989 cohort | 1.04<br>(0.729)<br>[5-40] | 0.088**<br>(0.038)<br>[19-55] | -0.028<br>(0.981)<br>[32-60] | -0.053*<br>(0.029)<br>[20-60] | 1.54***<br>(0.273)<br>[20-53] | 0.013<br>(0.021)<br>[35-55] |
| 1993 cohort | 1.19*<br>(0.716)<br>[10-40] | - | 0.139<br>(0.710)<br>[28-60] | - | 2.06*<br>(1.06)<br>[10-60] | - |

**Notes:** \*Significant at 10% level \*\*Significant at 5% level \*\*\*Significant at 1% level
The table shows the IV estimates of the effect of track attendance on years of schooling (YoS) and the log of the average monthly wage, using Model (3), for the three choice margins. Wage effects are not estimated for the 1993 cohort because of a lack of first stage power. Standard errors are between parentheses and are robust and corrected for clustering at the school level. Bandwidths are based on the cross-validation procedure and are between brackets.

### 5.2 Wages

Columns 2, 4 and 6 of Table 3 show the effects of track assignment on monthly wages. Results indicate that attending T2 vs. T1 at the margin leads to higher monthly wages of around 15% for the 1977 cohort (ages 36-42) and the 1983 cohort (ages 30-36) and around 9% for the 1989 cohort (ages 24-30). For T4 treatment, results show a positive effect of around 7% for the 1977 cohort. Wage estimates are positive but statistically insignificant for the 1983 and 1989 cohort. Results by year show that estimates for the 1989 cohort steadily increase across the time period 2001-2007 and are statistically significant in the more recent years. These estimates point to wage gains of around 4 to 5%. As such, it seems likely that are (small) wage gains for the 1989 cohort when they have built up a few years of labor market experience. This is in line with [13], who show that wages increase more sharply with experience in academic tracks. Estimates for the 1983 cohort are, however, consistently low. Comparing the 1977 and 1983 cohorts at the same age (2001 vs 2007 wages, respectively), the 1977 cohort still has considerably stronger wage gains (estimates are consistently around 7% in all years for the 1977 cohort). Hence, the difference clearly represents a cohort effect rather than a wage effect.

Estimates of the effect of track attendance on wages are statistically significant and negative for the T3 vs. T2 margin. This contrasts with the other two margins, where attending the higher track improves wages, and also with the lack of an effect on years of schooling. This suggests that marginal students are often pushed into a T3 track when they would be better off in a T2 track, from a lifetime earnings perspective. It is not surprising that this is not yet reflected in the years of schooling results as the T3 students attend a higher track and are eligible for higher levels in post-secondary education. One could therefore say that the lack of a positive effect for educational attainment already signals a high potential fo sorting too many students into T3 at the T2/T3 margin.

The question remains what concretely drives these negative wage effects. As mentioned before, the near-zero effects on average years of schooling hide some degree of substitution of *mbo* diplomas for more *hbo* diplomas but fewer post-secondary diplomas in general. It could be that the return to *mbo* diplomas is comparatively strong at this margin. Data on study choice might provide an additional explanation. We find that attendance of T3 over T2 at the margin leads to increases in study choice towards health and, especially, humanities, and a decrease in exact sciences and, especially, economics (see Appendix Table A2). Average wages are considerably higher in the latter compared to the former. Hence, study choice can explain part of these negative wage results. We can only speculate on the reason why these study choice patterns emerge, but it could be that being in a more demanding track and being ranked lower compared to classroom peers leads students to move away from studies that are perceived as having ‘more challenging’ curricula.

As mentioned before, the wage data are corrected for full-time equivalent of the job, exclude very low wages, and are topcoded at 20,000 euro’s per month. Wage results are similar when using uncorrected wages. Estimates for T2 vs. T1 treatment are somewhat larger. Similarly, we find that T2 vs. T1 treatment implies an increase in the full-time equivalent of the job and a (slight) increase in employment. We identify negative point estimates for FTE in case of T4 vs. T3 treatment, which is statistically significant in the 1983 cohort.

Comparing the IV estimates (Table 3) to the OLS estimates (Table 2), the results for the two higher choice margins are more favorable for attending the higher track in the OLS model. This likely reflects the positive bias in the ATE estimates due to students self-selecting into tracks. For T2 vs. T1 treatment, the positive wage estimates are higher for the IV model. As a strong negative bias in the ATE appears unlikely, this is likely driven by the local nature of IV estimates. The IV model estimates the effect of T2 vs. T1 for students that are induced to switch to a higher track when they have higher achievement levels. This could be students that are especially ambitious and they might also attend relatively better schools (they are more likely to attend schools with stronger entrance requirements which is likely to imply better peer quality). This could explain the relatively high local estimates in this particular case.

### 5.3 School achievement

Table 4 shows the effect of track assignment on school achievement in grade 9. While the focus of this study is on the long-run effects of track attendance, achievement could provide a potential mechanism towards such long-run effects. As explained in Section 4, the 9th grade test results in the 1989 and 1993 cohort contain a large share of (selective) missing values. To deal with this, we provide two imputation approaches. The first imputes missing tests from the relevant domain of the Entrance test. Taking into account that those that did not take the test might have developed especially poorly between grades 7 and 9, we subtract half a standard deviation from the imputed values in the second imputation approach. Comparing these alternative approaches provides an indication of the robustness of the results to the issue of missing values. Panel B of Table 4 shows results for the 1999 cohort, in which the share of missing tests is negligible. The data for this cohort also contains a problem-solving test. There are only two margins to estimate here, as tracks T1 and T2 were merged in 1999.

**Table 4:** IV estimates of the effect of track assignment on 9th grade test scores

| Panel A: 1989 and 1993 |  |  |  |  |  |  |
| --- | --- | --- | --- | --- | --- | --- |
|  | T2 vs. T1 |  | T3 vs. T2 |  | T4 vs. T3 |  |
|  | Language | Math | Language | Math | Language | Math |
| 1989 Base | 0.136<br>(0.239) | 0.0092<br>(0.286) | -0.020<br>(0.198) | 0.302*<br>(0.165) | 0.641***<br>(0.098) | 0.141<br>(0.096) |
| 1989 Impute I | 0.043<br>(0.123) | 0.131<br>(0.167) | -0.102<br>(0.118) | 0.072<br>(0.110) | 0.418***<br>(0.072) | 0.121*<br>(0.063) |
| 1989 Impute II | -0.091<br>(0.136) | -0.020<br>(0.179) | -0.154<br>(0.134) | 0.018<br>(0.132) | 0.434***<br>(0.087) | 0.153*<br>(0.079) |
|  | [5-40] | [5-40] | [20-50] | [20-50] | [20-53] | [20-53] |
| 1993 Base | 0.781**<br>(0.273) | -0.339<br>(0.329) | -0.0070<br>(0.266) | 0.0022<br>(0.152) | 0.836**<br>(0.361) | 0.436*<br>(0.230) |
| 1993 Impute I | 0.107<br>(0.122) | 0.247<br>(0.160) | -0.0088<br>(0.135) | -0.027<br>(0.085) | 0.512***<br>(0.218) | 0.090<br>(0.121) |
| 1993 Impute II | 0.108<br>(0.126) | 0.266*<br>(0.158) | -0.151<br>(0.170) | -0.035<br>(0.113) | 0.465*<br>(0.279) | 0.069<br>(0.186) |
|  | [10-40] | [10-40] | [20-60] | [20-60] | [10-60] | [10-60] |
| Panel B: 1999 |  |  |  |  |  |  |
|  | T3 vs. T1/T2 |  |  | T4 vs. T3 |  |  |
|  | Language | Math | PS | Language | Math | PS |
| Entrance test | 0.292**<br>(0.135) | 0.321<br>(0.206) | -0.122<br>(0.158) | 0.186*<br>(0.097) | 0.454*<br>(0.247) | 0.083<br>(0.166) |
|  | [10-60] | [18-60] | [15-60] | [20-60] | [30-51] | [30-60] |
| Cito test | 0.299**<br>(0.132) | 0.461*<br>(0.237) | -0.0024<br>(0.140) | 0.405*<br>(0.209) | 0.406*<br>(0.237) | 0.080<br>(0.082) |
|  | [515-542] | [510-539] | [520-550] | [510-547] | [534-548] | [520-550] |

**Notes:** \*Significant at 10% level \*\*Significant at 5% level \*\*\*Significant at 1% level
The table shows the IV estimates of the effect of track assignment on school achievement in grade 9, using Model (3). ‘Impute 1’ imputes missing scores from the same domain on the Entrance test. ‘Impute 2’ additionally subtracts 0.5 standard deviation from the imputed values. Panel B shows results for the 1999 cohort, in which T1 and T2 now form one merged track, separately for when using the Entrance Test (low stakes) or the Cito test (high stakes) as forcing variable. The new merged track is labeled as ‘T1/T2’. ‘PS’ refers to a problem-solving test. Standard errors are between parentheses and are robust and corrected for clustering at the school level. Bandwidths are based on the cross-validation procedure and are between brackets. The full range for the Cito test score is [501-550].

Panel A of Table 4 shows that results are indeed sensitive to imputation, but the positive treatment effect for T4 vs. T3 appears robust, especially for language. Positive effects at this margin are also identified for the 1999 cohort. In Panel A, there is no (robust) evidence for an effect on achievement at the other margins. In contrast, panel B provides positive effects at the lower margin for both language and math. The impact of track assignment on the problem-solving test is low and statistically insignificant for both margins. This could reflect that such skills are less driven by instruction and peer effects or, more generally, difficult to influence at later ages.

The positive results for language and math for the lower margin appear to contradict panel A, but these results are difficult to compare because of the merger of the two lowest tracks. This has effectively created a new margin. Moreover, the 1999 cohort constitutes the first cohort after the policy change and we could therefore also pick up on transition effects. We conclude that there appear to be positive achievement effects for T4 vs. T3 at the margin, while we do not find any achievement effects at the lower margins under the old system. It should be emphasized that these tests elicit *academic* achievement, while attending vocational tracks can potentially benefit students’ skills in non-academic disciplines. The data to test the impact of track attendance on vocational skills are not available.

The identified effect for T4 vs. T3 appears predominantly driven by peer effects. Adding average peer quality in class as a control leads to low and statistically insignificant estimates. The literature on peer effects suggests that an increase in peer quality of 1 standard deviation leads to an increase in individual achievement by around 0.40 of a standard deviation [20]. The difference in peer achievement between T3 and T4 is around 0.75 of a standard deviation, indicating the effect sizes are in line with what the literature predicts on the basis of peer effects.

The 1999 cohort also contains data for the high-stakes Cito test, which allows a comparison of using either test as forcing variable. Panel B of Table 4 shows that using either the Entrance Test or the Cito test leads to similar estimates. This result is in line with the observation from Figure A2 that the Cito score is not necessarily a stronger predictor of track assignment. Hence, having only a proxy for the true selection test does not appear to impact the estimates in a strong way.

These results indicate an improvement in cognitive skills from attending the higher track, at least at the higher margin. Track attendance can potentially also effect non-cognitive skills, which are also highly important for future earnings [21]. The data are relatively limited when it comes to such outcomes, e.g. measurement of Big Five personality skills are missing. Cohorts 1977, 1989 and 1993 do contain 9th grade measures for school enjoyment and ‘need for achievement’. The latter can be seen as a proxy for motivation, and has been shown to be highly predictive for educational outcomes [22]. Effects for school enjoyment are low, and only statistically significant for T4 vs. T3 treatment in 1989 (with positive sign). Effects for ‘need for achievement’ are positive for T2 vs. T1 treatment and strongly negative for T3 vs. T2 treatment in all cohorts. Hence, the long-run treatment effects at these margins could be partially driven by intermediate effects on non-cognitive skills.

### 5.4 Heterogeneity across background characteristics

We further analyze whether the effect of track assignment differs across observable characteristics (these results are not shown but available on request). For this purpose, we include an interaction between the attended track and specific background characteristics and instrument this variable with an interaction between the instrument and the specific characteristic. In general, the precision of the interaction estimates is low, especially for the T3 vs. T2 margin, but they reveal some interesting patterns. We first of all look at interactions with background characteristics. For gender, point estimates suggest weaker positive effects on years of schooling for women at the lower margin, but these are not statistically significant. We identify a statistically significantly stronger effect for women with respect to years of schooling at the higher margin in 1977, but not for any of the other cohorts. Point estimate with respect to wages are consistently low. We further estimate interactions with parental education. Conformity suggests that those with highly educated parents could be especially prone to be assigned to too demanding tracks. We do not find evidence for this. We identify slightly weaker positive effects for those with higher educated parents with respect to years of schooling at the higher margin, and also somewhat for wages (only for the 1989 cohort). All other interaction estimates are low and statistically insignificant. A potential explanation is that higher educated parents also have more means to support their child if it is struggling in a more demanding track.

Additionally, we estimate interactions between treatment and achievement indicators, namely the Entrance Test score and the teacher recommendation given at the end of primary school. These estimations provide insights into how effects differ by perceived ability, and in the extent to which the main estimates truly reflect the treatment effect for those at the margin (namely those with a mixed teacher recommendation). With respect to the Entrance Test score, we identify positive point estimates as one would expect given the depiction in the theoretical model, but these are generally low and only statistically significant with respect to years of schooling at the highest margin. Hence, we do not identify strong heterogeneity by achievement but this is likely the result of a lack of statistical power and the fact that students from a specific track can be concentrated within a relatively small range. In other words, a large part of the range of the functions portrayed in Fig 1 is not observed in reality.

Interactions with the teacher recommendation show similar results as for the Entrance Test. More importantly, the (summed) point estimates for those with a mixed recommendation are very close to the main estimates, across cohorts, outcomes and margins. In other words, the identified effect in the main model appears highly representative of the group with a teacher recommendation right at the margin.

### 5.5 Discussion

For students at the lowest and at the highest choice margin, being assigned to the higher track provides a return in the form of higher wages, but at the cost of time and resources spent on education. The wage gains can be a complete result of the increase in educational attainment, but other mechanisms can be in effect as well. As such, we cannot state that the wage gain represents the ‘return’ on the extra investment in schooling, but if one wants to assess whether individuals are (economically) better off when attending the higher track, years of education and wages represent the relevant costs and benefits of that choice. From that perspective, one extra year of education has a ‘return’ of around 10% on monthly wages for T2 vs. T1 treatment. For T4 vs. T3 treatment, the payoff is around 7% for the 1977 cohort, and negligible if we look at average wages for the 1983 and 1989 cohorts. If we take the more recent wage data for the 1989 cohort, the return for an extra year of education is around 3% at age 30.

The return to schooling is generally found to be around 8 or 9% in the literature; see, e.g., [23, 24, 25]. However, these averages hide a considerable amount of heterogeneity, and might not be representative of returns at the achievement margin. Returns to schooling at the margin of dropout are often found to be especially high [25]. This could also explain why our wage estimates are larger at the lowest margin. [10] estimate an average return to an extra year of college of 14%, but a marginal return that ranges from 1.5% to 8.5% (depending on the specific margin the policy change affects). The ‘return’ we identify for T4 treatment is in line with those effect sizes.

The results for the highest margin may be seen as especially low given the positive effects on achievement. However, earlier cited studies have shown that attendance of higher tracks often leads to short-term increases in achievement but no or weak labor market returns. Study choice may also provide an explanation for the low returns. For the T4 vs. T3 margin, we also find a relative increase in attendance of post-secondary studies with lower average wages, mainly towards humanities (see Appendix Table A2). This also likely explains the slight negative effect on the FTE of the job at the T4 vs. T3 margin, as the hours worked for those with majors in these areas tend to be lower. From an individual perspective, it remains inconclusive whether students around the margin are better off in the higher track. Many individuals do not pursue higher education even when expected returns are high, due to, e.g., income risk and psychic costs of studying [26]. Whether attending the higher track represents a net gain or a net loss for the marginal student ultimately depends on his or her utility function.

Such ambiguity does not apply to the T3 vs. T2 margin, as there is a strong negative wage effect, for the same average years of schooling. Hence, thresholds for this margin appear to be too lenient. This indicates that either parents and students are more likely to push for the higher track when close to this achievement margin, and/or that schools set more lenient entry requirements. The former could occur because completing T3 provides direct access to higher education and could therefore be seen as an especially crucial threshold. For schools, the incentives to set lower thresholds indeed appear comparatively high at this threshold. Tracks 3 and 4 typically fall within the same school while T2 and T3 do not. As such, schools are incentivized to set higher thresholds between T3 and T4 to increase average student quality within each track, while they are incentivized to set lower thresholds between T2 and T3 to attract more students. We lack the data to empirically assess these potential explanations.

It should be emphasized that, in this setting, ‘too lenient’ thresholds do not strictly imply that the threshold scores that schools set are, on average, too low. As there is leeway for parents with high aspirations for their children to still get into higher tracks also when they have relatively low scores, or to switch to another school with more lenient requirements, it can also be the case that such behavior pushes higher formal (though lenient) thresholds into lower effective thresholds. Hence, negative wage returns for attending T3 over T2 could potentially also be avoided by stricter adherence to thresholds, rather than increasing the thresholds as such. In any case, our results indicate to parents that pushing students into higher tracks around the margin is not necessarily beneficial for later-life outcomes.

Our main findings appear to contrast with those of [14], who find no long-run effect for either educational attainment or wages from attending the higher track. However, as stated before, they estimate treatment effects at a very different margin. The lack of any effect in their study does not imply that ability thresholds are placed ‘optimally’. For example, it can also reflect that underambitious allocation of younger students is canceled out by overambitious allocation of older students. Additionally, [14] suggest that their zero effects can be a consequence of the relatively high upward flexibility in the German system. The Dutch tracking system is comparatively more rigid with respect to upward mobility between tracks, which can also explain the difference in findings.

## 6 Robustness

We now assess the sensitivity of our results towards different specifications and robustness tests. Many of the tests we conduct are similar to those for the traditional RD design, as the identification threats are similar (although based on more lenient assumptions) compared to our non-linearity approach. As stated before, where the RDD assumes no discontinuity in other determinants, our design assumes linearity in other determinants across the specified range of the running variable. We critically assess this assumption by analyzing sensitivity to the inclusion of observable characteristics, to bandwidth choice and to excluding tracks that are not part of the relevant margin. The latter two tests also addresses the issue to what extent our design still estimates treatment effects at the achievement margin. Finally, we assess sensitivity of our results to how the non-linearity in treatment probability is exactly exploited.

### 6.1 Observable characteristics

The estimation approach assumes that any other determinants of the outcome variable are linearly related to achievement and thereby captured by the control function for the Entrance Test score. When the inclusion of control variables strongly changes the estimates this is a strong indication that this assumption is invalid. The main results presented before include the set of controls *X′*. Table 5 compares those to a model without controls. The changes in the estimates are all small. This result confirms descriptive statistics shown before that indicated that the relation between achievement and control variables is linear. Fig 7 has shown that some non-linearity is present at the higher end for wages in the 1983 cohort. The results from Table 5 show that this has no major impact on the estimates, as sensitivity in the 1983 cohort is not larger than in other cohorts. The underlying reason is that any such non-linearity is also reflected in the CV procedure for optimal bandwidths, which consequently suggests narrower bandwidths.

**Table 5:**
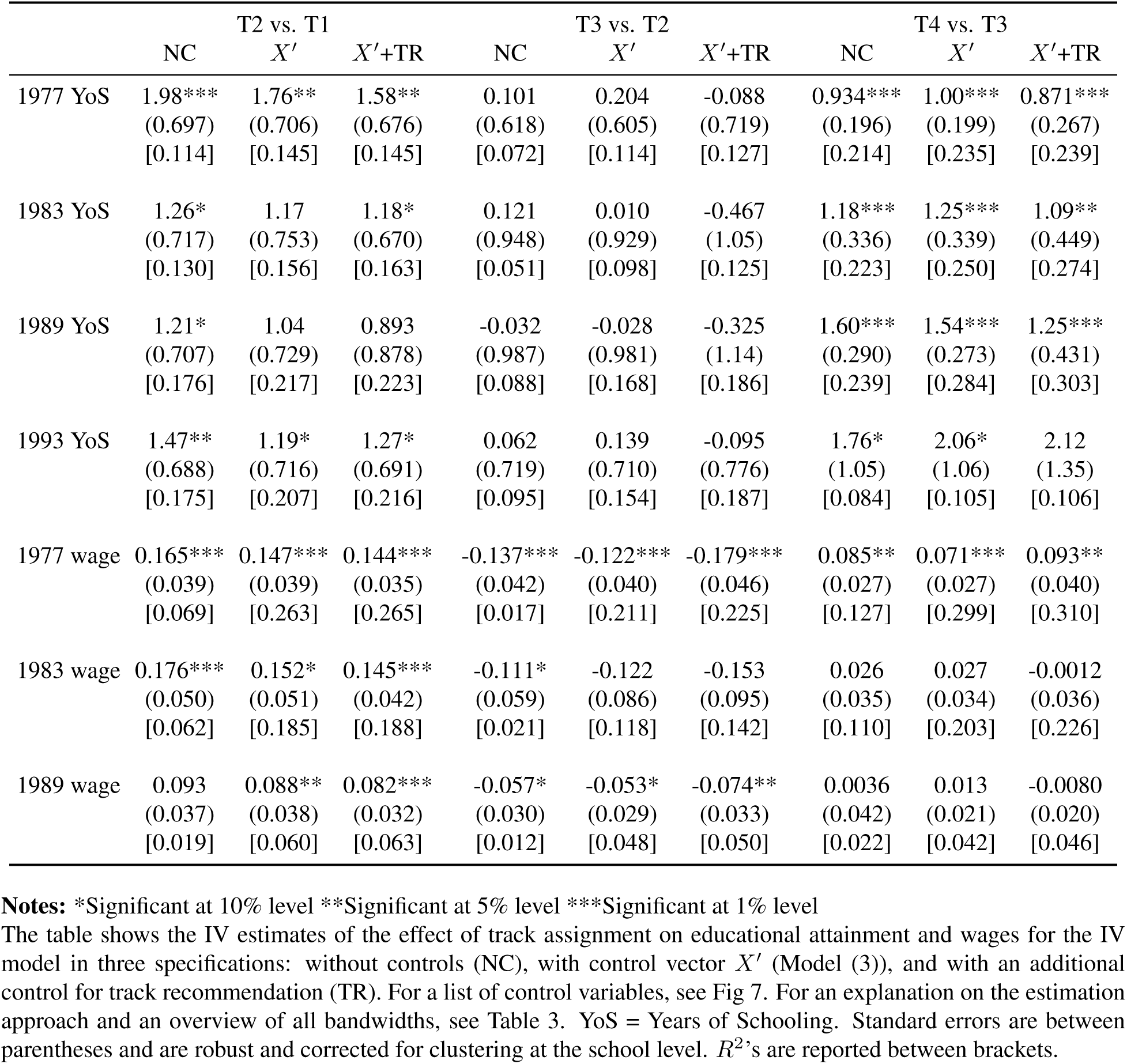
IV estimates of the effect of track assignment: with and without controls

Table 5 also shows estimates when we additionally include a control for the teacher recommendation that students receive at the end of primary school. Track recommendations correlate strongly with the Cito exit test, but can differ when teachers feel that the test does not accurately reflect student ability or potential. As such, the variable provides a valuable control for aspects that are important for future success but not fully captured by test scores, such as non-cognitive skills and motivation. If our estimates would strongly respond to controlling for the teacher recommendation, it would suggest that they are partly driven by non-linearity in such unobserved determinants across the control function. Addition of the teacher recommendation reduces first stage power somewhat, partly because the variable is not available for around 5% of the sample. Table 5 shows that the sensitivity to this additional control is very minimal.

As shown by [27], coefficient stability when adding controls should always be judged in relation to the explanatory power of those control variables. Table 5 also reports the *R*^2^ of the different specifications. The results show that control variables have substantial explanatory power, especially towards wages and especially for higher tracks, and that the teacher recommendation further adds to this. The fact that the latter explains outcomes also conditional on student achievement and background suggests that it indeed captures other types of skills than a test score. The sensitivity of the estimates is not higher in cases where the *R*^2^ increases more strongly. We cannot rule out that non-linearities in unobserved determinants of long-run outcomes bias our results. Nonetheless, the low sensitivity of the estimates to the inclusion of observed indicators lends further validity to our results. The high coefficient stability also in cases where the *R*^2^ increases substantially implicitly indicates that any potential selection on unobservables (conditional on *S_i_*) would have to be very strong to fully drive these estimates.

A more formal exercise to assess possible non-linearity in important observable characteristics is to conduct placebo tests that use constructed control vectors as outcome variables in Model 3. Results of this exercise are shown in Table A3 in the appendix. Only one of the placebo tests is statistically significant (at the 10% level). Given that we test this for 21 different hypotheses, these results support the assumption that important observable determinants are linearly related to our outcomes, and therefore are captured by the control function approach. We similarly obtain statistically insignificant estimates when the teacher recommendation is used as outcome.

### 6.2 Sensitivity to bandwidth

Our non-linearity model has been estimated after establishing optimal bandwidths, following the same approaches as in RDD designs. Table 6 shows how sensitive our results are to modifications in the bandwidth choice. The table shows results for years of schooling and wage estimates in the 1977 and 1983 cohorts, at all three margins (results for the 1989 and 1993 cohorts are not shown but available on request). The first column shows the results estimated with the bandwidth suggested by the CV procedure. The next columns show the effects of a range of changes in that bandwidth. Appendix Table A4 provides a full overview of bandwidths used in this exercise.

**Table 6:** Sensitivity of results to bandwidth

| T2 vs. T1 |  |  |  |  |  |  |  |  |
| --- | --- | --- | --- | --- | --- | --- | --- | --- |
|  | BL | LL- | LL+ | LL++ | UL- | UL- - | UL+ | UL++ |
| 1977 YoS | 1.76**<br>(0.706) | 1.84***<br>(0.694) | 1.87***<br>(0.729) | 1.79**<br>(0.743) | 2.81***<br>(0.942) | 2.25**<br>(1.01) | 1.51***<br>(0.445) | 1.11**<br>(0.323) |
| 1983 YoS | 1.17<br>(0.752) | 1.64***<br>(0.636) | 0.823<br>(1.06) | 3.33*<br>(1.98) | 2.63***<br>(0.938) | 2.32*<br>(1.36) | 2.15***<br>(0.642) | 1.82***<br>(0.621) |
| 1977 wage | 0.147***<br>(0.039) | 0.148***<br>(0.039) | 0.144***<br>(0.039) | 0.135***<br>(0.044) | 0.104**<br>(0.049) | 0.243***<br>(0.072) | 0.161***<br>(0.035) | 0.158***<br>(0.035) |
| 1983 wage | 0.152***<br>(0.051) | 0.138***<br>(0.050) | 0.139**<br>(0.062) | 0.186***<br>(0.082) | 0.121**<br>(0.056) | 0.133*<br>(0.073) | 0.150***<br>(0.050) | - |
| T3 vs. T2 |  |  |  |  |  |  |  |  |
|  | BL | LL- | LL- - | LL+ | LL++ | UL- | UL- - | UL- - - |
| 1977 YoS | 0.204<br>(0.605) | -0.626<br>(0.440) | -0.633*<br>(0.344) | -0.935<br>(0.992) | 0.134<br>(1.35) | 0.330<br>(0.614) | 1.05<br>(0.642) | 1.13<br>(0.807) |
| 1983 YoS | 0.010<br>(0.929) | -0.853<br>(0.602) | -1.37***<br>(0.476) | 1.33<br>(1.38) | 2.28<br>(1.74) | -0.326<br>(0.928) | -0.378<br>(0.928) | 0.036<br>(1.03) |
| 1977 wage | -0.122***<br>(0.040) | -0.130***<br>(0.036) | -0.138***<br>(0.035) | -0.152***<br>(0.049) | -0.140**<br>(0.065) | -0.124***<br>(0.040) | -0.090**<br>(0.042) | -0.064<br>(0.049) |
| 1983 wage | -0.122<br>(0.086) | -0.098*<br>(0.057) | -0.110**<br>(0.045) | -0.189<br>(0.154) | -0.078<br>(0.178) | -0.123<br>(0.086) | -0.116<br>(0.087) | -0.099<br>(0.090) |
| T4 vs. T3 |  |  |  |  |  |  |  |  |
|  | BL | LL- - | LL+ | LL++ | LL+++ | UL- | UL- - | UL+ |
| 1977 YoS | 1.00***<br>(0.199) | 1.01***<br>(0.194) | 1.14***<br>(0.211) | 1.14***<br>(0.235) | 1.25***<br>(0.283) | 1.11***<br>(0.231) | 0.592<br>(0.403) | 0.966***<br>(0.184) |
| 1983 YoS | 1.25***<br>(0.339) | 1.02***<br>(0.271) | 1.36***<br>(0.461) | 0.692<br>(0.739) | 0.509<br>(1.19) | 1.11***<br>(0.361) | 0.985**<br>(0.446) | - |
| 1977 wage | 0.071***<br>(0.026) | 0.068***<br>(0.026) | 0.079***<br>(0.028) | 0.070**<br>(0.032) | 0.084**<br>(0.038) | 0.085**<br>(0.033) | 0.099**<br>(0.049) | 0.075***<br>(0.024) |
| 1983 wage | 0.027<br>(0.034) | 0.040<br>(0.027) | -0.016<br>(0.046) | -0.011<br>(0.069) | 0.066<br>(0.104) | 0.0083<br>(0.036) | 0.050<br>(0.045) | - |

**Notes:** \*Significant at 10% level \*\*Significant at 5% level \*\*\*Significant at 1% level

Results are consistent across the different bandwidths. More narrow bandwidths naturally lead to less precise estimates and larger variability, but also in those cases the qualitative conclusions remain the same, and effect sizes are in a similar range. In fact, comparatively larger changes occur when extending the bandwidths, especially at the lower end, which is not surprising as these extensions are rejected by the cross-validation exercise. While descriptive statistics showed some non-linearity at the higher end for the 1983 cohort (see Figs 6 and 7), the results are not more sensitive for changes in the upper bandwidth limit there. Although very narrow bandwidths are not feasible within our model, the fact that results are consistent for relatively more narrow ranges strongly suggests that the large optimal bandwidths are not what drives our findings.

### 6.3 Sample composition

We have argued before that our approach still elicits treatment effects close to the margin, because the LATE puts more weight on those observations for which the first stage slopes are steeper. The consistency of results for more narrow bandwidths provides evidence in favour of this. Moreover, we can assess how sensitive results are to the exclusion of tracks that are not part of the relevant margin.

In the main approach, we have included at least one other track in addition to the two marginal tracks in all but one case, as these sample compositions are favoured by the CV exercise. Appendix Table A5 shows results for the two possible alternatives in each case. As all alternatives involve a different sample, optimal bandwidths are re-estimated in each case. The model that only includes the two tracks at the margin (‘A2’) is naturally less precise, but the differences in the point estimates are not large and not consistent in one direction. Moreover, all main conclusions remain the same. Hence, the inclusion of individuals in the sample that are not part of the tracks at the choice margin does not affect our results. These extra observations mainly help in obtaining more precise estimates, but do not affect the point estimates of the LATE. The fact that even sample specifications that are rejected by our CV procedure produce results similar to our main estimates lends additional validity to the findings.

### 6.4 Smoothing out the conditional mean function

The main estimation approach exploits any changes that occur in treatment probability across *S_i_*. A potential problem is that this also incorporates variation in treatment probability that is the result of noise. When a set of students with a specific test score happens to be a ‘good draw’ by chance, this leads to both a higher probability of assignment to the higher track and likely better long-run outcomes as well. Coefficients would then be biased in favour of attending the higher track. We estimate an alternative approach in which we ‘smooth out’ any such volatility by predicting treatment by the Entrance test, assuming a logistic relation. These results are shown in Appendix Table A6. Estimates are naturally less precise in this approach, partly because we exclude noise but also because we might exclude any other variation that is not fitted by the assumed logistic functional form. This is mainly an issue for estimation of educational attainment effects for the medium margin. Hence, the result for this particular treatment effect remains somewhat inconclusive, and we cannot fully exclude that there is some effect of track assignment on years of schooling for this margin. Overall, point estimates In Table A6 are very similar to the main results. The small differences that do occur indeed point to slightly lower estimates in the logistic approach, as we would expect, but these are very minor and all the main conclusions are still upheld. Hence, volatility in treatment probability that is caused by noise does not drive our estimation results.

## 7 Conclusion

This study has assessed the long-run effect of secondary school track assignment for students who at the margin of the required achievement level for a specific track. The empirical approach relies on a design that exploits non-linearity in the relation between treatment probability and achievement, while assuming that achievement is linearly related to other determinants of the outcome variable. The fact that track assignment is based on (implicit) achievement thresholds leads to a situation in which increases in the probability of attending a higher track are not proportional to increases in ability or potential. Descriptive statistics and various robustness analyses lend support to the assumption that the non-linearity in the relationship with achievement is exclusive to the probability of track assignment. Our results indicate that students in the Netherlands obtain higher educational attainment and higher wages when attending the higher track, for the choice margin between Tracks 1 and 2 and the choice margin between Tracks 3 and 4. The returns in the labor market are around 15% for the lower margin and between 3% and 7% for the higher margin, at the expense of around 1.5 additional years of schooling. Estimates for the choice margin between the two middle tracks are low and statistically insignificant with respect to educational attainment (although imprecisely estimated), and negative with respect to wages, indicating a wage loss of around 12% from attending the higher track.

Several potential mechanisms can drive these treatment effects. Attending higher tracks directly provides access to higher levels of post-secondary education, which subsequently is linked to higher expected earnings in later life. Additionally, different tracks imply different curricula that teach different types of skills, as well as differences in peer and school quality. We find robust evidence of positive effects of higher track attendance on school achievement for the higher margin, which is suggestive of true learning effects. In light of the positive effects for achievement and educational attainment, the labor market returns at the higher margin appear to be low. Additionally, labor market returns are negative for the middle margin, where there is no effect on educational attainment. These patterns appear to be explained at least partly by study choice. Attending a higher track leads to more frequent sorting into study majors with lower future earnings. This is suggestive evidence that the more challenging track in secondary school leads students to select ‘less challenging’ post-secondary educational paths. It also relates to recent findings by [28] that a lower rank in class (which attending a higher track directly induces) leads to lower investment in human capital, conditional on ability. We also identify negative treatment effects on motivation at the middle margin, which could contribute to this pattern of study choice. Further disentangling the different potential mechanisms that are behind the effects of track assignment is an interesting avenue for future research.

The results from this study highlight that, while higher tracks are associated with higher wages, the students who are at the achievement margin of a track do not necessarily obtain such strong wage gains. In our study, wage returns depend strongly on the margin we are looking at: high at the lower margin, positive but low compared to educational investments at the higher margin, and negative at the middle margin. We can only speculate on the underlying reasons for these differences. As discussed before, different considerations of students and their parents and of schools can influence the location of the thresholds. Parental aspirations for the educational attainment of their children are known to be high and this can lead parents to push children into too demanding tracks. Our pattern of results could reflect that such parental overconfidence would be less prominent among the low-achieving children. Additionally, the negative results for the middle margin could partly accrue due to the structure of Dutch schools. As the majority of T3 schools also offer T4 but not T2, there is a comparatively stronger incentive for schools to lower the threshold for T3 to attract more students. This variation in treatment effects across margins also makes it difficult to project the sign and size of track assignment effects in other countries. Replication of our analysis for those countries (possibly with a similar empirical approach) would be needed to assess the external validity of our findings.

For the lowest and highest choice margin, wage gains come at the expense of extra investment in education. Whether this is perceived as a gain or a loss for the individual depends on his or her utility function. The particular approach developed in this paper does not answer what is the optimal choice from the perspective of society. A lower threshold can negatively affect the untreated, because peer quality is reduced on both sides of the threshold. On the other hand, higher educational attainment could induce externalities for society in terms of reduced crime rates or productivity spillovers. The social welfare implications of different achievement thresholds for track assignment can be an interesting avenue for future research, for example by looking at exogenous variation in assignment over time induced by policy changes. At the same time, such policy changes cannot be used to assess the implications of choosing the higher over the lower track for the individual student at the margin, because they reassign a whole segment of the distribution and thereby also change the nature of each track. Hence, one has to rely on another source of exogenous variation.

Ideally, the effects of track assignment would be estimated by exploiting a strong discontinuity in explicit threshold scores for track eligibility. Despite the comparatively strong reliance on achievement in track assignment in the Dutch system, the traditional RD design was not feasible, even in the case where the official running variable is available. The identification approach therefore relies on the assumption that unobservable determinants of long-run outcomes are linearly related to achievement, which we cannot formally prove. While relying on stronger assumptions, the conditional mean approach presented in this study can provide an alternative approach for empirical studies with similar data designs.

## Acknowledgments

The authors would like to thank Thomas Dohmen, Andries de Grip, Tyas Prevoo, Trudie Schils, the participants of the EALE conference in Bonn (2012) and seminar participants at Maastricht University (2012, 2017) and KU Leuven (2015) for their valuable comments.

